# The Common Marmoset as a Translational Model for Longitudinal Studies of Cognitive Aging and Individual Vulnerability to Decline

**DOI:** 10.1101/2024.08.22.609213

**Authors:** Casey R. Vanderlip, Payton A. Asch, Courtney Glavis-Bloom

## Abstract

In humans, cognitive aging is highly variable, with some individuals experiencing decline while others remain stable, and different cognitive domains exhibiting uneven vulnerability to aging. The neural mechanisms driving this intra- and inter-individual variability are not fully understood, making longitudinal studies in translational models essential for elucidating the timelines and processes involved. The common marmoset (*Callithrix jacchus*), a short-lived nonhuman primate, offers an unprecedented opportunity to conduct longitudinal investigations of aging and age-related disease over a condensed time frame, in a highly translatable animal model. The potential of the marmoset as a model for cognitive aging is indisputable, but a comprehensive cognitive battery tailored for longitudinal aging studies has not yet been developed, applied, or validated. This represents a critical missing piece for evaluating the marmoset as a model and understanding the extent to which marmoset cognitive aging mirrors the patterns found in humans, including whether marmosets have individual variability in their vulnerability to age-related cognitive decline. To address this, we developed a comprehensive touchscreen-based neuropsychological test battery for marmosets (MarmoCog), targeting five cognitive domains: working memory, stimulus-reward association learning, cognitive flexibility, motor speed, and motivation. We tested a large cohort of marmosets, ranging from young adults to geriatrics, over several years. We found significant variability in cognitive aging, with the greatest decline occurring in domains dependent on the prefrontal cortex and hippocampus. Additionally, we observed significant inter-individual variability in vulnerability to age-related cognitive decline: some marmosets declined across multiple domains, others in just one, and some showed no decline at all. This pattern mirrors human cognitive aging, solidifies the marmoset as an advantageous model for age-related cognitive decline, and provides a strong foundation for identifying the neural mechanisms involved.

## INTRODUCTION

Age-related cognitive decline is a key feature of many neurodegenerative disorders, with abilities in some cognitive domains declining earlier and more severely than others, due to their reliance on neural circuits more vulnerable to the effects of aging (Salthouse, 2009; Hara et al., 2012; Stark et al., 2013). Even within vulnerable domains, age-related cognitive decline is not ubiquitous; only select individuals exhibit decline. Despite significant efforts, the neural mechanisms driving variable domain and inter-individual susceptibility to cognitive decline remain poorly understood. Longitudinal approaches provide dynamic insights that are crucial for identifying the onset, phenotype, progression, and severity of cognitive decline that are not possible with cross-sectional studies alone. This information is vital to both determining and modifying the neural mechanisms driving age-related cognitive decline.

Longitudinal studies in humans are impractical due to the decades of observation necessary to examine the extent of our long lifespan. Therefore, to address scientific questions that are best answered using longitudinal approaches, we must turn to animal models like monkeys, which have shorter lifespans and significant translational relevance to humans. Monkeys share similar physiology and neuroanatomy with humans, including a well-developed prefrontal cortex, and spontaneously develop age-related and Alzheimer’s disease-associated neuropathological changes, including beta-amyloid deposition, hyperphosphorylation of tau, reduced hippocampal neurogenesis, and synaptic dysfunction (Morrison and Baxter, 2012; Ngwenya et al., 2015; Arnsten et al., 2021; Glavis-Bloom et al., 2023; Perez-Cruz and Rodriguez-Callejas, 2023). Furthermore, monkeys can perform complex cognitive tasks similar to those used in human clinical diagnoses and exhibit domain-specific, age-related cognitive impairment, with a subset of monkeys resistant to age-related cognitive decline (Rapp and Amaral, 1989; Gray and Barnes, 2019; Sadoun et al., 2019; De Castro and Girard, 2021; Glavis-Bloom et al., 2022; Baxter et al., 2023; Vanderlip et al., 2023, 2024a). Among monkeys, macaques are the most widely used, but their lifespan, which exceeds 30 years, severely limits the practicality of longitudinal studies of aging processes.

Recently, the common marmoset (*Callithrix jacchus*) has emerged as an advantageous monkey model for neuroscience research. As a nonhuman primate, the marmoset maintains all of the same value of macaques, with a substantial additional benefit: a short lifespan. Marmosets are the shortest-lived anthropoid primates, living on average for 12 years, and considered aged at seven (Tardif et al., 2011). This short lifespan offers an unprecedented opportunity to conduct longitudinal investigations into aging and age-related disease over a condensed time frame, in a highly translatable animal model.

The vast potential of the marmoset as a model for studying aging is indisputable, but an important question remains unanswered: Do aging marmosets exhibit domain-specific, individual variability in vulnerability to cognitive decline? Here, we addressed this question by developing a comprehensive touchscreen-based neuropsychological test battery for marmosets (MarmoCog) that covers five cognitive domains: working memory, stimulus-reward association learning, cognitive flexibility, motor speed, and motivation. We used MarmoCog to assess cognitive performance of a large cohort of marmosets ranging in age from young adult through geriatric, and conducted repeated testing over several years. We show that performance in some cognitive domains is more vulnerable to decline than others. Additionally, individual marmosets exhibited different patterns of cognitive aging: some declined across multiple domains, some in only one, and others showed no decline at all. This work firmly establishes the marmoset as an advantageous model of cognitive aging and sets the stage for investigation of the neurobiological underpinnings of individual vulnerability to age-related cognitive decline.

## METHODS

### Subjects

Nineteen (10 female, 9 male) common marmosets (*Callithrix jacchus*), initially 2.84 to 16.73 years of age, were administered MarmoCog twice per year for several years. The marmosets were singly or pair-housed in enriched cages with visual and auditory access to other marmosets. Pairs were separated during cognitive testing and were not food or water restricted. All experiments complied with the Institutional Animal Care and Use Committee of the Salk Institute for Biological Studies and conformed to NIH guidelines.

### Equipment

Marmoset home cages were custom-built to include a testing chamber located in the top corner of the cage, with a 10.4-inch infrared touch screen testing station (Lafayette Instrument Company, Lafayette, IN) mounted in place. Access to this testing chamber was restricted except during cognitive testing sessions during which marmosets could freely enter and exit through a small doorway. They earned liquid rewards, such as apple juice, dispensed into a sink centered beneath the touchscreen via a peristaltic pump. Cognitive task-specific “masks” were secured in front of the screen. These masks were made of plastic and had cutouts to delineate potential stimulus locations. Animal Behavior Environment Test Cognition software (Lafayette Instrument Company, Lafayette, IN) was used to program and administer all cognitive tasks, and also recorded task events such as stimuli displays and touches.

### Cognitive Testing

Cognitive tests were administered 3-5 days per week for up to 3 hours. Daily testing sessions ended when three hours had elapsed, 20mL of reward was earned, or the task-specific completion criteria had been met. All stimuli were simple black-and-white shapes displayed on a black background. Detailed descriptions of all tasks have been previously published and are described briefly below (Glavis-Bloom et al., 2022; Vanderlip et al., 2023).

#### Working Memory: Delayed Recognition Span Task (DRST) and Delayed Non-match-to-Sample (DNMS)

Working memory capacity was assessed using the Delayed Recognition Span Task (DRST). Marmosets initiated each trial by touching a blue square stimulus in the center of the screen. Following this, a randomly chosen stimulus from a set of 400 images was displayed in one of nine possible locations on the screen. Upon selection, a small liquid reward was given. After a two-second delay with a blank screen, a two-alternative forced choice was presented between the original stimulus in its initial location and a new stimulus in a different location. Accuracy on this beginning portion of each DRST trial was also analyzed separately because it constitutes a trial of the commonly used Delayed Non-match-to-Sample (DNMS) task. If the marmoset selected the new stimulus, a correct response was logged, a reward was dispensed, and another two-second delay followed. Subsequently, the first two stimuli reappeared in their original locations along with a third, new stimulus in a new location. The marmoset was rewarded again for choosing the new stimulus. This process continued, with novel stimuli added after each delay, until the trial ended in one of three ways: the marmoset 1.) made nine correct selections in a row, 2.) failed to make a selection within 12-seconds (an omission), or 3.) made an incorrect response (chose a non-novel stimulus). In cases of omission or incorrect response, no reward was given, and a five-second time-out was implemented before a new trial started. For each trial, the number of correctly selected stimuli before termination was recorded as the “Final Span Length” and used as the primary dependent variable. The reward volume increased with the trial’s difficulty to maintain engagement.

#### Stimulus-Reward Association Learning: Visual Discrimination (VD)

Stimulus-reward association learning was assessed with a Visual Discrimination (VD) task. After a trial was initiated, a pair of black-and-white stimuli appeared side-by-side in randomly assigned positions. One stimulus in the pair was predetermined to be the correct one, the other incorrect. If the correct stimulus was selected, the marmoset earned a small liquid reward. If the incorrect stimulus was selected or if no choice was made within 12-seconds (an omission), no reward was given and a five-second time-out preceded the five-second inter-trial interval. Marmosets continued to receive two-alternative forced choice trials with the same pair of stimuli until they achieved 90% accuracy (18 correct out of 20 consecutive trials). The primary dependent variable was the number of errors to criterion for each Problem.

#### Cognitive Flexibility: Serial Reversal (SR)

Cognitive flexibility was evaluated using a Serial Reversal (SR) task. After completing a single VD Problem as described above, without any indication to the marmoset, the reward contingencies were reversed. Thus, the previously rewarded stimulus became unrewarded, and the previously unrewarded stimulus became rewarded (i.e., Reversal 1). The marmosets learned the new reward contingencies through trial and error until achieving the 90% performance criterion. Then, the reward contingencies were reversed again, restoring the original conditions (i.e., Reversal 2). The primary dependent variable was the number of errors to criterion on each Reversal.

#### Motor Speed: Reaction Time Task (RTT)

Motor speed was measured using a Reaction Time Task (RTT). After the trial was initiated, a target appeared in one of nine pseudo-randomly chosen locations. When the marmoset touched the target, they received a small liquid reward. The primary dependent variable was the elapsed time from target onset to selection on each trial. Each test session ended after 108 trials were completed or three hours elapsed, whichever came first.

#### Motivation: Progressive Ratio Task (PRT)

Motivation was measured using a Progressive Ratio Task (PRT). Each trial began with a stimulus in the center of the screen. Marmosets were rewarded for touching the stimulus according to a progressive-ratio schedule where response requirements increase during the session. Specific increments were: 1, 2, 3, 4, 5, 6, 7, 8; followed by increments of 2: 10, 12, 14, 16, 18, 20, 22, 24; and increments of 4: 28, 32, 36, 40, 44, 48, 52, 56; and so on. Once each response requirement was met, the stimulus was removed, a reward dispensed, and then the stimulus reappeared. Test sessions ended after one hour. The primary dependent variable was the number of rewards earned in the one-hour session.

### Data Cleaning

Since the goal of this study was to assess the effects of aging on longitudinal cognitive performance, it was necessary to filter the data to isolate the effects of aging from confounding factors related to task acquisition and initial learning (Glavis-Bloom et al., 2022; Vanderlip et al., 2023). Therefore, the data included in this study is from after the marmosets were trained to a high proficiency level on the cognitive tasks. After proficiency was demonstrated, marmosets were tested on each of the cognitive tasks approximately every six months for several years. Each of these testing instances is referred to as a “time point”.

For the DRST, the first time point was defined by the first 300 trials after marmosets reached asymptotic performance (Glavis-Bloom et al., 2022). Some marmosets performed the DRST continuously for up to three years, with subsequent time points identified every six months, each including up to 300 trials. After transitioning to other cognitive tasks, DRST performance continued to be assessed every six months. Omitted trials and anticipatory responses (less than 500ms in the same location as the trial initiation stimulus) were excluded from analyses. Since DNMS trials were embedded in DRST trials, the time points were identically defined. Omitted trials with one or two stimuli displayed and anticipatory responses were excluded from analyses.

Initial acquisition of the VD task consisted of learning the stimulus-reward associations of six sequential Problems. We demonstrated that, for the first two Problems, aging was associated with impaired performance (Vanderlip et al., 2023). Therefore, for the first time point of this study, to isolate age-related decline from learning deficits, we included data from only the last three of these Problems. Subsequently, marmosets completed three Problems per time point. Omitted trials were not included in the calculations of errors to criterion.

During the initial acquisition of the SR task, marmosets completed one VD Problem followed by five serial reversals of that single pair of stimuli (Vanderlip et al., 2023). To avoid confounding learning effects, data from only the last two of these reversals were included in the first time point. For all subsequent time points, marmosets completed two reversals on each of the three learned VD Problems. Omitted trials were not included in the calculations of errors to criterion.

For RTT, marmosets initially completed ten sessions of maximally 108 trials each. To control for initial learning of the task, and the fact that not all marmosets completed the full 108 trials offered in each session, we included the last 432 trials in the first time point. For subsequent time points, all marmosets were tested until they completed 432 trials, with a maximum of 216 trials in a single daily session. Trials that were omitted or trials where the target stimulus was presented in the same location as the trial initiation stimulus were excluded from analyses.

Initial testing of the PRT consisted of between four and eleven one-hour sessions. To standardize across animals, data from the last four of these initial sessions were included in the first time point. Subsequent time points each consisted of four one-hour sessions.

### Statistical Analyses

All statistical analyses and data visualization were performed in Python. Nonparametric tests were used throughout due to the non-normal distribution of the data. To determine whether an individual marmoset’s performance declined on a given task over time, we used Mann-Kendall tests. The Mann-Kendall test is a statistical approach to identify significant trends over time from longitudinal data and avoids the potential for bias inherent in experimenter-defined performance cutoffs. Marmosets were identified as declining on a task when their Mann-Kendall test was significant (p < 0.05) with a negative tau value. Only data from marmosets with three or more time points on a given task were statistically analyzed. To visualize individual marmoset performance over time, we used linear or logistic regressions for continuous and binary data, respectively. We used the same approaches to analyze and visualize performance as a function of age across animals on each task. To evaluate potential associations between performance on different tasks, we created a correlation matrix using Spearman’s correlation coefficients and applied a Bonferroni correction to adjust for multiple comparisons.

## RESULTS

In this study, we conducted a comprehensive multi-year investigation of cognitive aging in marmosets. We characterized cognitive aging trajectories in five distinct cognitive domains: working memory, stimulus-reward association learning, cognitive flexibility, motor speed, and motivation. By evaluating performance on these domains over the marmoset lifespan, we evaluated the patterns and extent of age-related cognitive decline across and within individuals.

### Working Memory

To investigate whether marmosets experience longitudinal cognitive decline in working memory, we assessed the DRST and DNMS (Figure 1A) performance of 14 marmosets approximately every six months for up to five years. The monkeys ranged in age from 2.96 to 16.30 years at the beginning of testing (Figure 1B).

**Figure 1.**
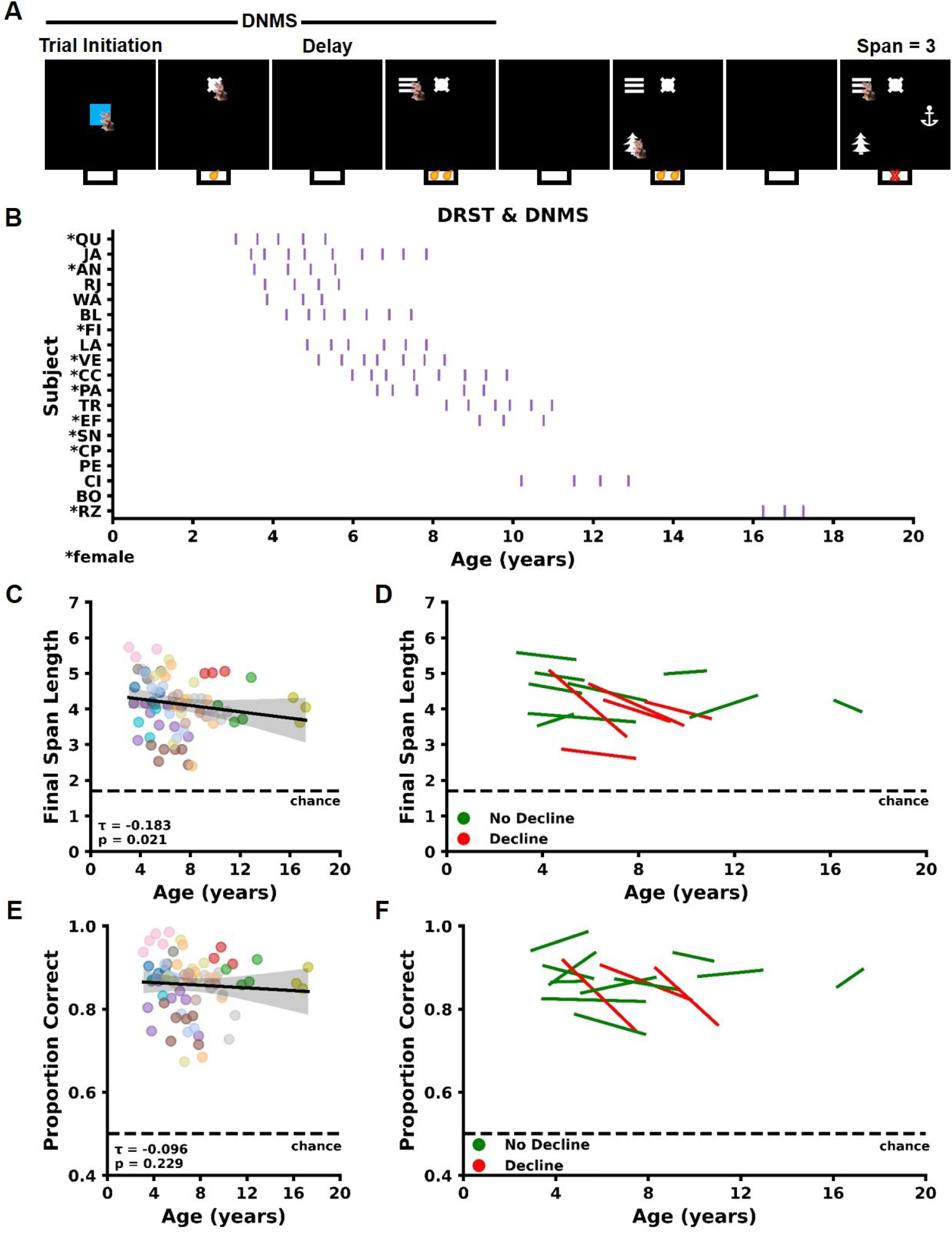
Longitudinal changes in working memory. A) Schematic of the Delayed Recognition Span Task (DRST) with the embedded Delayed Nonmatch-to-Sample (DNMS) task. B) Chart depicting the age of each testing time point for each marmoset. Each vertical line on the chart represents one testing time point. C) Linear regression across individuals shows that increasing age is associated with reduced Final Span Length across the cohort. Each dot represents mean performance for a single individual for a single time point. The dots are colored by individual marmoset. Gray shading indicates 95% confidence interval determined by linear regression model. D) Linear regression of DRST performance over time for each individual marmoset. A subset of five marmosets exhibits significant decline (red lines) on the DRST over time, while other marmosets did not decline (green). E) Increasing age is not associated with changes in DNMS performance across the cohort. Dots, colors, and shading as in C. F) Linear regression of DNMS performance over time for each individual marmoset. The performance of three marmosets significantly declined on the DNMS task (red lines). All three marmosets who declined on DNMS also declined on the DRST.

#### Delayed-Recognition Span Task (DRST)

As in prior work, we used Final Span Length, defined as the number of stimuli the marmoset correctly identified as novel on each trial, as the primary measure to assess DRST performance (Herndon et al., 1997; Moss et al., 1997; Killiany et al., 2000; Moore et al., 2017; Glavis-Bloom et al., 2022; Vanderlip et al., 2024a). We first investigated whether performance on the DRST declined with age. Consistent with prior research, we found a negative trend in Final Span Length with increasing age, indicating a decline in working memory capacity (Figure 1C; Mann-Kendall τ = −0.18, p = 0.021). To determine whether this decline was ubiquitous across all marmosets or driven by a subset, we conducted Mann-Kendall tests for each marmoset individually. Our analysis revealed that five of the 14 marmosets (36%) showed significant negative performance trends, indicating a decline in working memory capacity with age (Table 1; Figure 1D). The remaining nine marmosets showed no significant negative trends, indicating no decline in working memory capacity despite advancing age (Figure 1D). The marmosets with declining working memory varied in age, including middle-aged individuals, while some older marmosets remained stable, highlighting significant inter-individual differences in cognitive aging among marmosets.

**Table 1.**
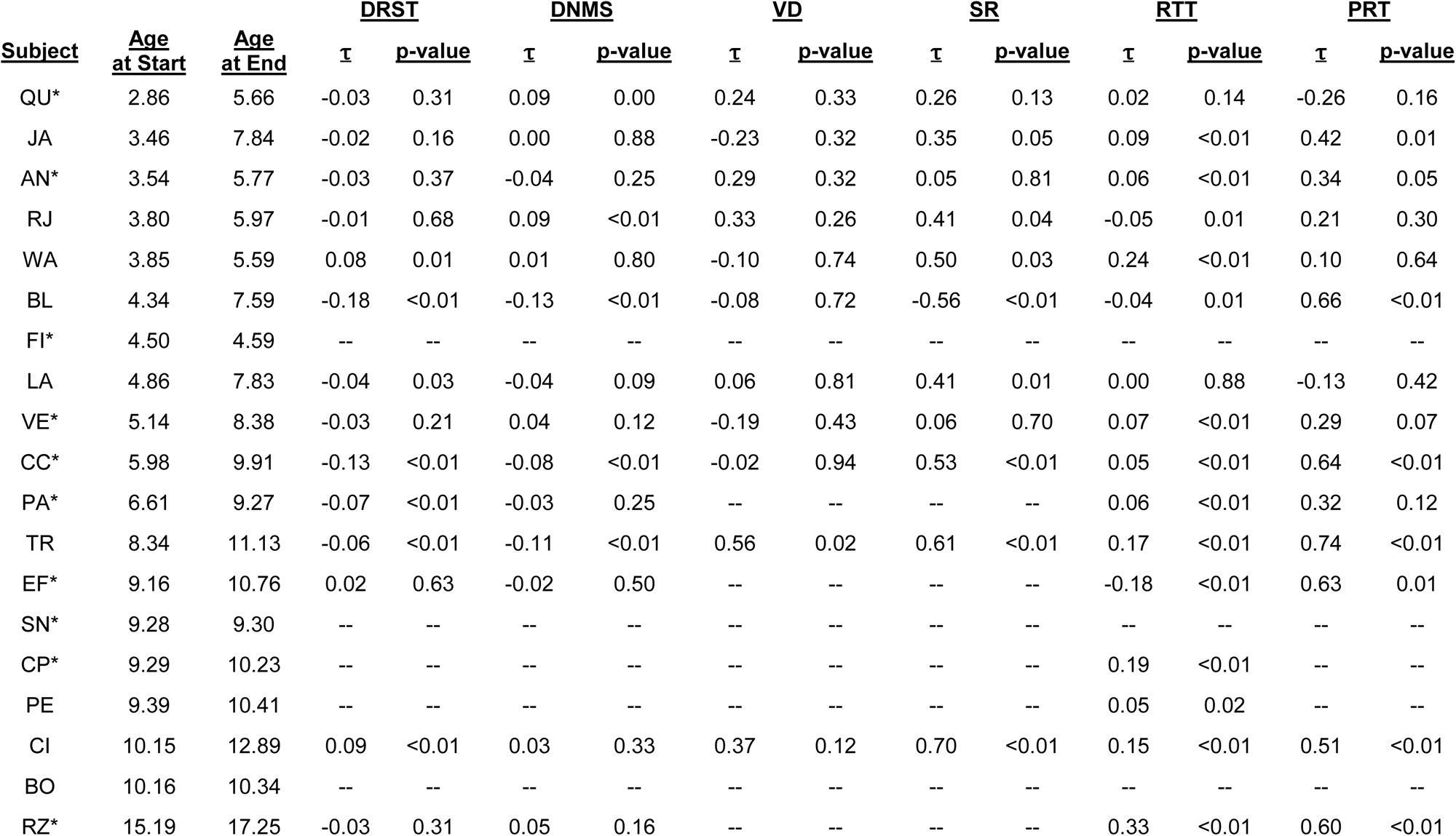
Statistical results from Mann-Kendall tests for trends in performance over time.

#### Delayed Non-match-to-Sample Task (DNMS)

We next analyzed the portion of each DRST trial that constituted a trial of the well-known DNMS task. Across the cohort, no age-related decline in accuracy on the DNMS was observed (Figure 1E; Mann-Kendall τ = −0.096, p = 0.229). However, Mann-Kendall tests to examine individual performance over time identified three marmosets whose accuracy decreased on the DNMS task (Table 1; Figure 1F). All three of these marmosets also declined on the DRST, while two other marmosets declined on the DRST but not on the DNMS. This suggests that, though both of these tasks assess working memory, performance on these two tasks is not equivalent; the DRST is more sensitive to decline, likely because it is more difficult than the DNMS.

### Stimulus-Reward Association Learning

We investigated longitudinal cognitive decline in marmosets’ stimulus-reward association learning using the VD task (Figure 2A). We evaluated 18 marmosets approximately every six months for up to three years, with initial ages ranging from 3.35 to 16.71 years (Figure 2B). First, we analyzed how performance changed with age across all marmosets and found no significant age-related changes in errors to criterion (Figure 2D; Mann-Kendall τ = −0.043, p = 0.655). Next, we determined that, of the 11 marmosets who completed at least three time points of the VD task, none had significant negative performance trends (Table 1; Figure 2E). This stable ability in stimulus-reward association learning with age is consistent with findings in humans and macaques (Rapp, 1990; Lai et al., 1995; Cox et al., 2008), and suggests that the capacity for stimulus-reward association learning is preserved across different species and is unaffected by aging.

**Figure 2.**
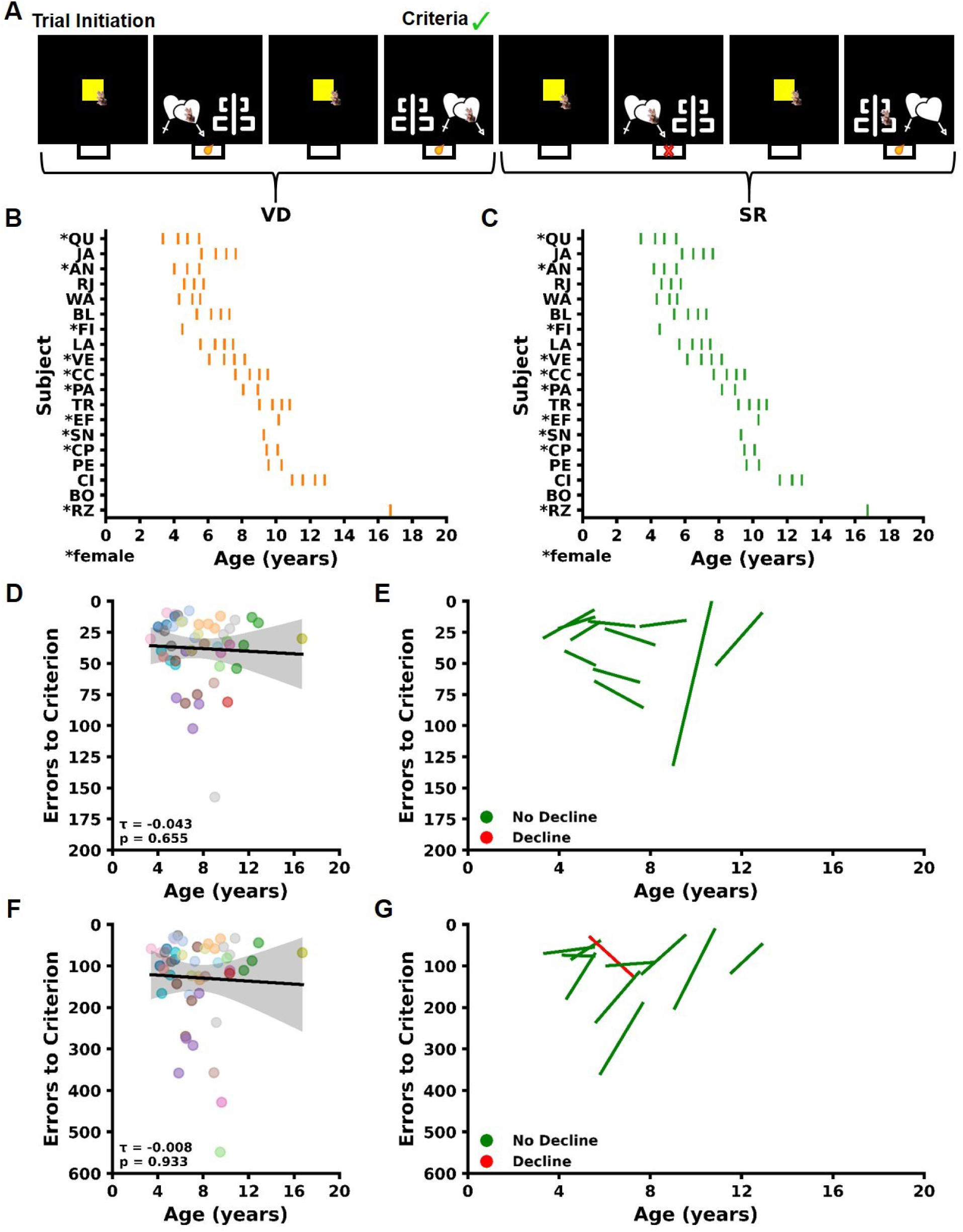
Age-related changes in stimulus-reward association learning and cognitive flexibility. A) Schematic of the Visual Discrimination (VD) and Serial Reversal (SR) tasks. Charts depicting the age of each testing time point for each marmoset on the B) VD and C) SR tasks. Each vertical line represents one testing time point. Linear regression across the cohort of marmosets shows that D) stimulus-reward association learning and F) cognitive flexibility remain stable with increasing age, when analyzed as a group. Each dot represents mean performance for a single individual for a single time point. The dots are colored by individual marmoset. Gray shading indicates a 95% confidence interval determined by the linear regression models. E) Plot of linear regression lines for each individual marmoset on the VD task. Marmosets with no significant decline are plotted in green. No marmosets significantly declined over time, confirming what was seen in the group regression that stimulus-reward association learning does not decline with advancing age. G) Linear regression plots for each individual marmoset on the SR task. Line color indicates no decline (green) or significant decline (red). One marmoset exhibited significant decline in cognitive flexibility with advancing age.

### Cognitive Flexibility

Longitudinal cognitive flexibility performance was assessed using the SR task. We evaluated the performance of 18 marmosets approximately every six months for up to three years, with initial ages ranging from 3.37 to 16.73 years (Figure 2B). First, we examined if SR performance declined with age and found no significant increase in the number of errors to criterion (Figure 2F; Mann-Kendall τ = −0.008, p = 0.933). Next, we conducted Mann-Kendall tests on data from marmosets that completed the task at least three times to determine individual performance trends. Our analysis showed that only one of the 11 marmosets (9%) exhibited a significant increase in the number of errors to criterion over time, indicating a decline in cognitive flexibility (Table 1; Figure 2G). The remaining 10 marmosets did not exhibit declining cognitive flexibility over the timeframe tested, despite advancing age. The one marmoset showing decline in cognitive flexibility also declined on the DRST which assesses working memory capacity. This multi-domain pattern of decline is suggestive of broader cognitive decline in this animal.

### Motor Speed

We investigated longitudinal cognitive decline in marmosets’ motor speed using the RTT (Figure 3A). We evaluated 18 marmosets approximately every six months for up to three years, with initial ages ranging from 2.85 to 15.20 years (Figure 3C). Consistent with previous research, we found that increasing age was associated with increased reaction times, indicative of motor slowing (Figure 3E; Mann-Kendall τ = −0.314, p < 0.0001). Next, we used Mann-Kendall tests to evaluate trends in motor speed over time for each marmoset. Among the 16 marmosets with at least three time points, three (20%) showed significantly increased reaction times suggesting motor speed decline (Table 1; Figure 3F). Notably, two of these marmosets were middle-aged, while one was aged, demonstrating that motor speed decline can occur at different life stages in different individuals.

**Figure 3.**
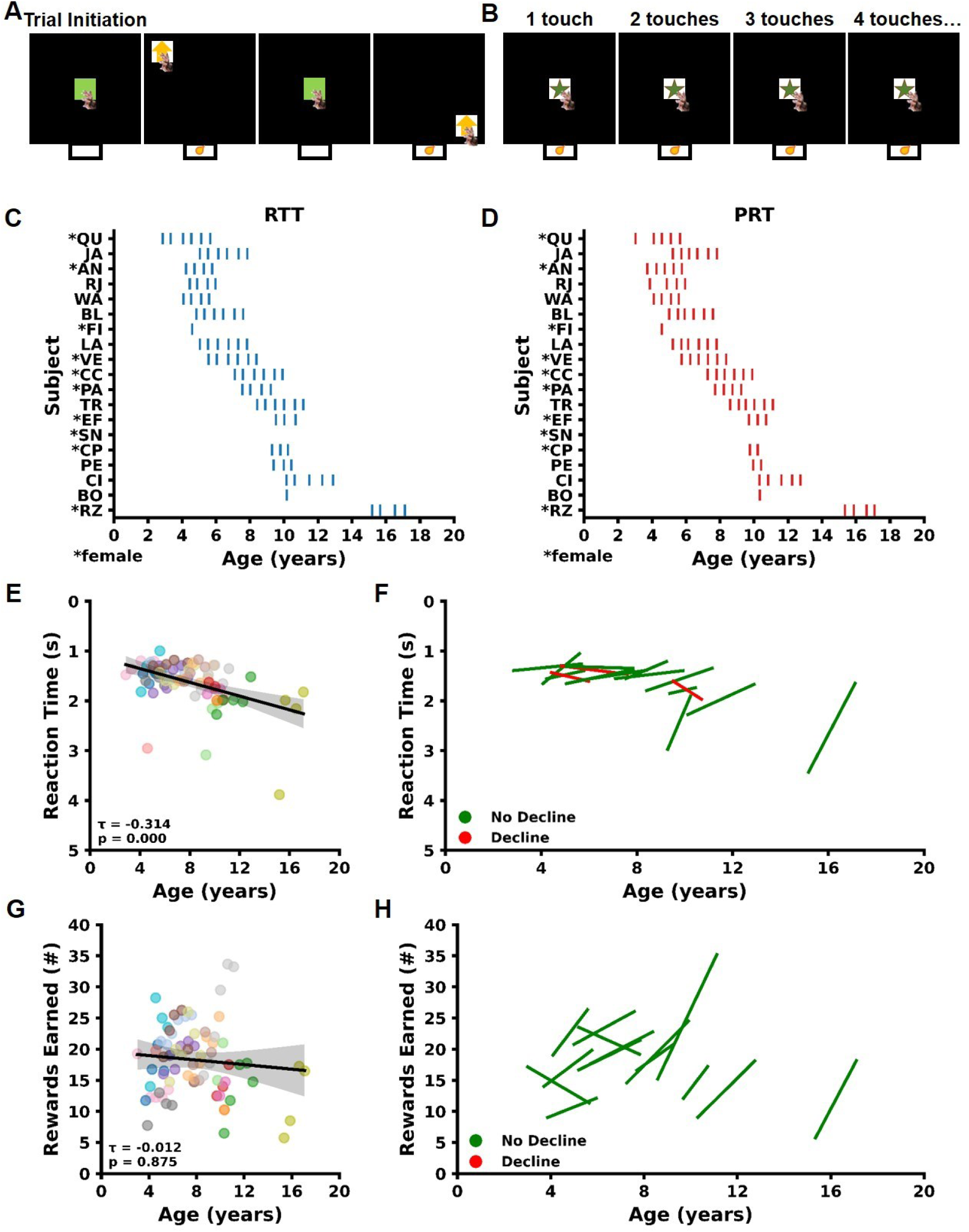
Age-related changes in motor speed and motivation. Schematics of the A) Reaction Time Task (RTT) and B) Progressive Ratio Task. Charts depicting the age of each testing time point for each marmoset on the C) RTT and D) PRT. Each vertical line represents one testing time point. E) Linear regression across the cohort of marmosets shows that increasing age is associated with motor slowing. Each dot represents mean performance for a single individual for a single time point. The dots are colored by individual marmoset. Gray shading indicates a 95% confidence interval determined by the linear regression models. F) Linear regression plots for each individual marmoset on the RTT. A subset of three marmosets showed significant motor slowing over time (red lines), while others did not (green lines). G) Linear regression across the cohort of marmosets shows that increasing age is not associated with changes in motivation. Dots, colors, and shading as in E. H) Linear regression plots for each individual on the PRT show that no marmosets exhibited significant decline in performance over time (green lines).

### Motivation

To investigate motivation trajectories in marmosets, we assessed their performance on the PRT. We evaluated the motivation of 18 marmosets approximately every six months for up to three years, with initial ages ranging from 3.01 to 15.37 years (Figure 3D). Across these animals, we found that age was not significantly associated with changes in motivation (Figure 3G; Mann-Kendall τ = −0.012, p = 0.875). We used Mann-Kendall tests to evaluate individual trends in motivation over time for 14 marmosets that completed the PRT at least three times. These analyses found that none of the 14 marmosets showed significant decline in motivation over the testing period (Table 1; Figure 3H).

### Intra- and Inter-Individual Patterns of Cognitive Decline

We systematically administered MarmoCog to 19 marmosets to assess performance in five separate cognitive domains over several years (Figure 4A). We observed multiple patterns of cognitive aging, with both intra- and inter-individual variability (Figure 4B). The most widespread decline was observed in the working memory domain on the DRST, where five marmosets exhibited significant declines in performance. Fewer instances of declining performance were observed on the other five tasks (DNMS, SR, RTT, PRT, and SD). Specifically, three marmosets declined on the secondary working memory task (DNMS), one marmoset declined on the SR task that measures cognitive flexibility, and three marmosets showed slowed motor speed on the RTT. In contrast, no marmosets showed decline in stimulus-reward association learning on the VD task, or reduced motivation on the PRT. Most frequently, marmosets exhibited decline in a single domain, indicating domain-specific cognitive aging. However, we found that working memory decline varied in its degree. Specifically, three marmosets were impaired on both working memory tasks (DNMS and DRST), while two others exhibited selective impairment only on the more complex DRST. However, one marmoset exhibited widespread cognitive decline with declining performance on four out of the six tasks. Together, these patterns suggest individual variability in the extent and nature of cognitive decline among aging marmosets.

**Figure 4.**
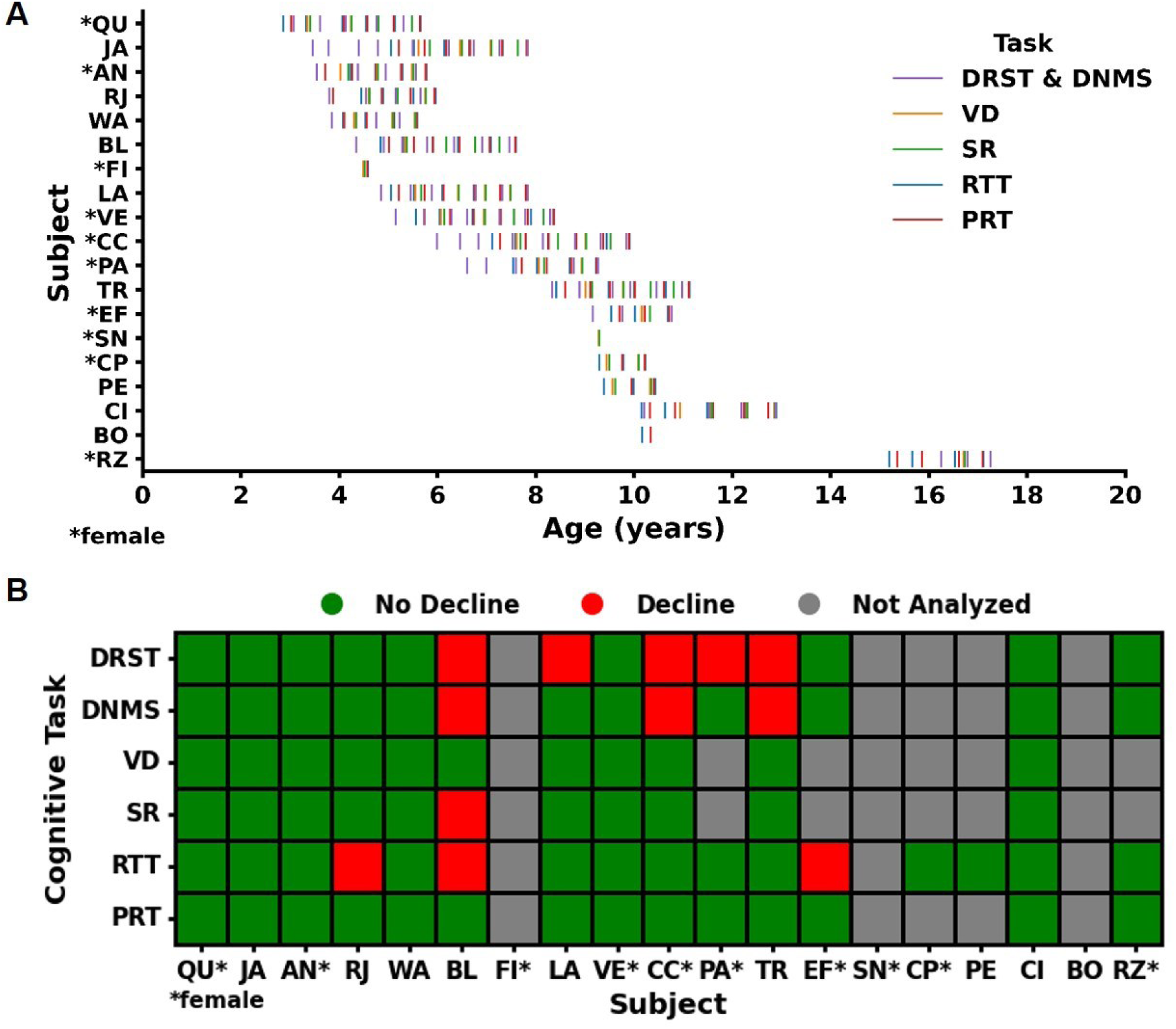
Longitudinal cognitive aging trajectories across domains. A) Chart depicting, with vertical lines, the age of each marmoset at each testing time point for each of the cognitive tasks. B) Heat map showing individual variability in cognitive decline patterns. One marmoset exhibited widespread cognitive decline across four cognitive tasks, others had significant declines in performance on the two tasks measuring the working memory domain (DNMS and DRST), some declined in a single domain, and others’ performance remained stable across all tasks.

### Correlational Analysis of Cognitive Task Performance

To determine whether performance on any of the six cognitive tasks was interrelated, we conducted a Spearman’s correlation analysis on data from the first time point of each task (Table 2). As expected, performance on the two working memory tasks (DNMS, DRST) were highly correlated (rs = 0.824, p = 0.004). Additionally, there was a significant positive correlation between performance on the cognitive flexibility task (SR) and each of the working memory tasks (DNMS: rs = 0.763, p = 0.023; DRST: rs = 0.741, p = 0.037). These significant correlations between tasks that assess different cognitive domains likely reflects their shared dependence on prefrontal cortical function. Importantly, we found no significant correlations between performance on either the motor speed (RTT) or motivation (PRT) tasks and any others (all ps > 0.1). This lack of associations strongly suggests that neither motor speed nor motivation contributes significantly to cognitive decline in other domains. Together, these results underscore the specificity of each cognitive task and highlight the independence of the domains they assess.

**Table 2.** Results from Spearman’s correlation analysis between tasks. * indicates p<0.05.

|  | <u>DRST</u> | <u>DNMS</u> | <u>VD</u> | <u>SR</u> | <u>RTT</u> | <u>PRT</u> |
| --- | --- | --- | --- | --- | --- | --- |
| <u>DRST</u> | 1.00 | 0.82* | 0.50 | 0.74* | 0.42 | 0.13 |
| <u>DNMS</u> | 0.82* | 1.00 | 0.58 | 0.76* | 0.38 | -0.18 |
| <u>VD</u> | 0.50 | 0.58 | 1.00 | 0.61 | 0.59 | 0.02 |
| <u>SR</u> | 0.74* | 0.76* | 0.61 | 1.00 | 0.38 | -0.07 |
| <u>RTT</u> | 0.42 | 0.38 | 0.59 | 0.38 | 1.00 | 0.57 |
| <u>PRT</u> | 0.13 | -0.18 | 0.02 | -0.07 | 0.57 | 1.00 |

## DISCUSSION

This study is the most extensive investigation of cognitive aging in the marmoset model to-date, both in terms of the multiple cognitive domains assessed, and the multi-year time frame of the assessments. By characterizing the cognitive aging trajectories of individual marmosets over most of their adult lives, we addressed critical questions about the marmoset’s validity as a model for aging, cognitive decline, and diseases for which these are a risk factor and symptom, respectively. We show that performance in cognitive domains requiring the prefrontal cortex and hippocampus is most vulnerable to age-related decline. Additionally, we found inter-individual variability in vulnerability to age-related cognitive decline, with some marmosets showing widespread decline across several domains, while others maintained largely stable performance. This pattern mirrors that of humans and macaques, solidifies the marmoset as an advantageous model for age-related cognitive decline, and lays a strong foundation for identification of the neural mechanisms underlying this decline.

### Development of MarmoCog: a cognitive battery for use in marmosets

Cognitive batteries are essential tools for measuring brain function across multiple cognitive domains and for identifying individuals with cognitive impairment. While batteries exist for humans and macaques, none does explicitly for marmosets (Weed et al., 1999; Hedden et al., 2012; Donohue et al., 2014; Baxter et al., 2023; Vanderlip et al., 2024b). The work of a few labs, taken together, demonstrates that marmosets are capable of performing different cognitive tasks (Spinelli et al., 2004; Sadoun et al., 2019; Glavis-Bloom et al., 2022; Vanderlip et al., 2023; Murai et al., 2024). However, in these studies, the subjects overwhelmingly were young, the tasks few, the majority of study designs cross-sectional, and some failed to detect age-related impairments seen robustly in humans and macaques (Sadoun et al., 2019; Rothwell et al., 2022; Murai et al., 2024). Furthermore, no study to-date has identified whether cognitive aging in marmosets is ubiquitous or whether there is inter-individual variability that is consistent with patterns known in humans.

In this study, we developed MarmoCog, a comprehensive cognitive battery spanning five domains: working memory, stimulus-reward association learning, cognitive flexibility, motor speed, and motivation. We demonstrated that marmosets of all ages can perform each of the tasks in MarmoCog. Moreover, marmosets performed the tasks voluntarily, without any food or water restriction, using home cage-mounted touch screens during short daily sessions, across their lifespan. This approach enables robust, high-throughput, no-stress, longitudinal cognitive testing. Furthermore, MarmoCog has unparalleled sensitivity, as demonstrated by our ability to identify individual marmosets with single, multi-domain, or no cognitive decline.

### Age-related decline on tasks dependent on the prefrontal cortex and hippocampus

Age-related neurobiological changes do not occur uniformly throughout the brain (Rapp and Amaral, 1989; Morrison and Baxter, 2012; Glavis-Bloom et al., 2023). Consequently, in humans and macaques, performance declines with age on tasks that engage the most vulnerable regions, such as the prefrontal cortex and hippocampus. To assess if this pattern holds true in the marmoset model, we longitudinally assessed performance on three tasks that critically depend on these most-vulnerable regions.

The DRST is a complex working memory task dependent on both the prefrontal cortex and hippocampus. Older adults perform worse on this task compared to younger adults, with impairments worsening in neurodegenerative conditions (Satler et al., 2015). Similar age-related impairments have been observed in marmosets and macaques in cross-sectional studies (Moss et al., 1997; Glavis-Bloom et al., 2022; Vanderlip et al., 2024a). Here, we examined marmoset working memory performance across more than half of their adult lifespan.

We found that approximately one-third of the marmosets tested (‘BL’, ‘LA’, ‘CC’, ‘PA’, ‘TR’) exhibited significant working memory decline on the DRST. Notably, three of these marmosets (‘BL’, ‘CC’, ‘TR’) also showed significant decline on the DNMS task, which is incorporated into the DRST but places less demand on working memory. Therefore, all marmosets with DNMS decline also declined on the DRST, while others declined only on the DRST. We hypothesize that these two patterns of working memory decline reflect different degrees of impairment. The marmosets with deficits on both tasks likely experience more severe working memory decline, as they are impaired even on the simpler DNMS task. Those with decline limited to the DRST may have a lesser degree of impairment, and it remains to be seen whether they will exhibit further decline as they age.

Cognitive flexibility, dependent on the prefrontal cortex, also declines with age in humans, macaques, and marmosets, as shown in cross-sectional studies (Rhodes, 2004; Moore et al., 2006; Head et al., 2009; Upright and Baxter, 2021; Gray et al., 2023; Vanderlip et al., 2023). However, the only longitudinal study in marmosets (Rothwell et al., 2022) found no age-related impairments, raising questions about the extent of such decline. In our longitudinal study, we also did not observe group-level impairment in cognitive flexibility when analyzing the data collectively. This may be due to most marmosets improving over time on the reversal task, masking previously identified deficits from our cross-sectional work (Vanderlip et al., 2023). However, by analyzing individual performance, we identified one marmoset (‘BL’) with clear cognitive flexibility decline, highlighting the importance of individual-level analysis in the assessment of cognitive decline.

This marmoset also showed decline on both working memory tasks, supporting the idea that age-related changes in the prefrontal cortex affect both working memory and cognitive flexibility. This is further supported by the significant correlation between performance on these tasks. Interestingly, other marmosets that showed decline in working memory did not decline in cognitive flexibility. Since the DRST engages both the prefrontal cortex and hippocampus (Beason-Held et al., 1999; Bor et al., 2006; Jeneson et al., 2010), this could indicate age-related hippocampal changes, such as hyperexcitability or microstructural changes, leading to reduced memory capacity (Yassa et al., 2011; Bakker et al., 2012; Venkatesh et al., 2020). Alternatively, different prefrontal circuits might underlie these functions, degrading independently or sequentially. This open question underscores the need for longitudinal studies across multiple cognitive domains to fully resolve the timeline of age-related brain changes.

### Age related changes in motor speed and motivation in marmosets

Accounting for potential confounds is critical for accurately identifying cognitive decline. Previously, we found significant motor slowing and decreased motivation with age in marmosets (Glavis-Bloom et al., 2022). Critically, there was no association between these age-related changes and cognitive impairments in cross-sectional, single time point evaluations (Glavis-Bloom et al., 2022). Here, we again observed group-level age-related motor slowing, with three individuals (‘RJ’, ‘BL’, ‘EF’), showing significant decline. Two of these marmosets showed no decline in other domains, while one (‘BL’) also declined in working memory and cognitive flexibility. Motor speed did not correlate with performance in these other domains, suggesting it did not account for these declines. Additionally, motivation did not decrease with age; in fact, most individuals showed increased motivation over time. The lack of correlation between motor speed, motivation, and cognitive performance reinforces the validity of MarmoCog for identifying individual vulnerability to age-related, domain-specific, cognitive decline.

## Conclusion

This multi-year longitudinal study represents the most comprehensive investigation of cognitive aging in marmosets to date. The development and subsequent longitudinal application of MarmoCog over most of the adult lifespan revealed domain-specific vulnerabilities and individual differences in marmoset cognitive aging patterns that mirror the human condition. These results both validate MarmoCog as a robust, high-throughput, sensitive, tool, and validate the marmoset as a highly valuable model for aging, cognitive decline, and the diseases for which these are a risk factor and symptom, respectively. Together, this work paves the way for future studies to elucidate the neural underpinnings of cognitive aging, identification of biomarkers that predict future risk of cognitive decline, and development of targeted interventions to mitigate this decline.

## Acknowledgments

This research was supported by an AHA-Allen Initiative in Brain Health and Cognitive Impairment award made jointly through the American Heart Association and The Paul G. Allen Frontiers Group: 19PABH134610000AHA, National Institutes of Health grants 1R21AG068967-01 and P51OD010425, grants from the Larry L. Hillblom Foundation and the Don and Lorraine Freeberg Foundation, and the Fiona and Sanjay Jha Chair in Neuroscience. We thank Katie Williams for assistance in the care of the marmosets and technical support.

## REFERENCES

Arnsten AFT, Datta D, Preuss TM (2021) Studies of aging nonhuman primates illuminate the etiology of early-stage Alzheimer’s-like neuropathology: An evolutionary perspective. Am J Primatol 83:e23254.

Bakker A, Krauss GL, Albert MS, Speck CL, Jones LR, Stark CE, Yassa MA, Bassett SS, Shelton AL, Gallagher M (2012) Reduction of Hippocampal Hyperactivity Improves Cognition in Amnestic Mild Cognitive Impairment. Neuron 74:467–474.

Baxter MG, Roberts MT, Roberts JA, Rapp PR (2023) Neuropsychology of cognitive aging in rhesus monkeys. Neurobiol Aging 130:40–49.

Beason-Held LL, Rosene DL, Killiany RJ, Moss MB (1999) Hippocampal formation lesions produce memory impairment in the rhesus monkey. Hippocampus 9:562–574.

Bor D, Duncan J, Lee ACH, Parr A, Owen AM (2006) Frontal lobe involvement in spatial span: Converging studies of normal and impaired function. Neuropsychologia 44:229–237.

Cox KM, Aizenstein HJ, Fiez JA (2008) Striatal outcome processing in healthy aging. Cogn Affect Behav Neurosci 8:304–317.

De Castro V, Girard P (2021) Location and temporal memory of objects declines in aged marmosets (Callithrix jacchus). Sci Rep 11.

Donohue MC, Sperling RA, Salmon DP, Rentz DM, Raman R, Thomas RG, Weiner M, Aisen PS, Australian Imaging, Biomarkers, and Lifestyle Flagship Study of Ageing, Alzheimer’s Disease Neuroimaging Initiative, Alzheimer’s Disease Cooperative Study (2014) The preclinical Alzheimer cognitive composite: measuring amyloid-related decline. JAMA Neurol 71:961–970.

Glavis-Bloom C, Vanderlip CR, Reynolds JH (2022) Age-Related Learning and Working Memory Impairment in the Common Marmoset. J Neurosci Off J Soc Neurosci 42:8870–8880.

Glavis-Bloom C, Vanderlip CR, Weiser Novak S, Kuwajima M, Kirk L, Harris KM, Manor U, Reynolds JH (2023) Violation of the ultrastructural size principle in the dorsolateral prefrontal cortex underlies working memory impairment in the aged common marmoset (Callithrix jacchus). Front Aging Neurosci 15:1146245.

Gray DT, Barnes CA (2019) Experiments in macaque monkeys provide critical insights into age-associated changes in cognitive and sensory function. Proc Natl Acad Sci 116:26247–26254.

Gray DT, Khattab S, Meltzer J, McDermott K, Schwyhart R, Sinakevitch I, Härtig W, Barnes CA (2023) Retrosplenial cortex microglia and perineuronal net densities are associated with memory impairment in aged rhesus macaques. Cereb Cortex 33:4626–4644.

Hara Y, Rapp PR, Morrison JH (2012) Neuronal and morphological bases of cognitive decline in aged rhesus monkeys. Age 34:1051–1073.

Head D, Kennedy KM, Rodrigue KM, Raz N (2009) Age differences in perseveration: Cognitive and neuroanatomical mediators of performance on the Wisconsin Card Sorting Test. Neuropsychologia 47:1200–1203.

Hedden T, Mormino EC, Amariglio RE, Younger AP, Schultz AP, Becker JA, Buckner RL, Johnson KA, Sperling RA, Rentz DM (2012) Cognitive profile of amyloid burden and white matter hyperintensities in cognitively normal older adults. J Neurosci Off J Soc Neurosci 32:16233–16242.

Herndon JG, Moss MB, Rosene DL, Killiany RJ (1997) Patterns of cognitive decline in aged rhesus monkeys. Behav Brain Res 87:25–34.

Jeneson A, Mauldin KN, Squire LR (2010) Intact working memory for relational information after medial temporal lobe damage. J Neurosci Off J Soc Neurosci 30:13624–13629.

Killiany RJ, Moss MB, Rosene DL, Herndon J (2000) Recognition memory function in early senescent rhesus monkeys. Psychobiology 28:45–56.

Lai ZC, Moss MB, Killiany RJ, Rosene DL, Herndon JG (1995) Executive system dysfunction in the aged monkey: Spatial and object reversal learning. Neurobiol Aging 16:947–954.

Moore TL, Bowley B, Shultz P, Calderazzo S, Shobin E, Killiany RJ, Rosene DL, Moss MB (2017) Chronic curcumin treatment improves spatial working memory but not recognition memory in middle-aged rhesus monkeys. GeroScience 39:571–584.

Moore TL, Killiany RJ, Herndon JG, Rosene DL, Moss MB (2006) Executive system dysfunction occurs as early as middle-age in the rhesus monkey. Neurobiol Aging 27:1484–1493.

Morrison JH, Baxter MG (2012) The ageing cortical synapse: Hallmarks and implications for cognitive decline. Nat Rev Neurosci 13:240–250.

Moss MB, Killiany RJ, Lai ZC, Rosene DL, Herndon JG (1997) Recognition memory span in rhesus monkeys of advanced age. Neurobiol Aging 18:13–19.

Murai T, Bailey L, Schultz L, Mongeau L, DeSana A, Silva AC, Roberts AC, Sukoff Rizzo SJ (2024) Improving preclinical to clinical translation of cognitive function for aging-related disorders: the utility of comprehensive touchscreen testing batteries in common marmosets. Cogn Affect Behav Neurosci 24:325–348.

Ngwenya LB, Heyworth NC, Shwe Y, Moore TL, Rosene DL (2015) Age-related changes in dentate gyrus cell numbers, neurogenesis, and associations with cognitive impairments in the rhesus monkey. Front Syst Neurosci 9 Available at: /pmc/articles/PMC4500920/?report=abstract [Accessed July 30, 2020].

Perez-Cruz C, Rodriguez-Callejas J de D (2023) The common marmoset as a model of neurodegeneration. Trends Neurosci 46:394–409.

Rapp PR (1990) Visual discrimination and reversal learning in the aged monkey (Macaca mulatta). Behav Neurosci 104:876–884.

Rapp PR, Amaral DG (1989) Evidence for task-dependent memory dysfunction in the aged monkey. J Neurosci Off J Soc Neurosci 9:3568–3576.

Rhodes MG (2004) Age-related differences in performance on the Wisconsin card sorting test: A meta-analytic review. Psychol Aging 19:482–494.

Rothwell ES, Workman KP, Wang D, Lacreuse A (2022) Sex differences in cognitive aging: a 4-year longitudinal study in marmosets. Neurobiol Aging 109:88–99.

Sadoun A, Rosito M, Fonta C, Girard P (2019) Key periods of cognitive decline in a nonhuman primate model of cognitive aging, the common marmoset (Callithrix jacchus). Neurobiol Aging 74:1–14.

Salthouse TA (2009) When does age-related cognitive decline begin? Neurobiol Aging 30:507–514.

Satler C, Belham FS, Garcia A, Tomaz C, Tavares MCH (2015) Computerized spatial delayed recognition span task: A specific tool to assess visuospatial working memory. Front Aging Neurosci 7:1–9.

Spinelli S, Pennanen L, Dettling AC, Feldon J, Higgins GA, Pryce CR (2004) Performance of the marmoset monkey on computerized tasks of attention and working memory. Cogn Brain Res 19:123–137.

Stark SM, Yassa MA, Lacy JW, Stark CEL (2013) A task to assess behavioral pattern separation (BPS) in humans: Data from healthy aging and mild cognitive impairment. Neuropsychologia 51:2442–2449.

Tardif SD, Mansfield KG, Ratnam R, Ross CN, Ziegler TE (2011) The Marmoset as a Model of Aging and Age-Related Diseases. ILAR J 52:54–65.

Upright NA, Baxter MG (2021) Prefrontal cortex and cognitive aging in macaque monkeys. Am J Primatol 83:e23250.

Vanderlip CR, Asch PA, Reynolds JH, Glavis-Bloom C (2023) Domain-specific cognitive impairment reflects prefrontal dysfunction in aged common marmosets. eNeuro Available at: https://www.eneuro.org/content/early/2023/08/07/ENEURO.0187-23.2023 [Accessed August 8, 2023].

Vanderlip CR, Jutras ML, Asch PA, Zhu SY, Lerma MN, Buffalo EA, Glavis-Bloom C (2024a) Parallel patterns of cognitive aging in marmosets and macaques. :2024.07.22.604411 Available at: https://www.biorxiv.org/content/10.1101/2024.07.22.604411v1 [Accessed August 22, 2024].

Vanderlip CR, Stark CEL, Initiative ADN (2024b) Digital cognitive assessments as low-burden markers for predicting future cognitive decline and tau accumulation across the Alzheimer’s spectrum. :2024.05.23.595638 Available at: https://www.biorxiv.org/content/10.1101/2024.05.23.595638v1 [Accessed July 9, 2024].

Venkatesh A, Stark SM, Stark CEL, Bennett IJ (2020) Age- and memory-related differences in hippocampal gray matter integrity are better captured by NODDI compared to single-tensor diffusion imaging. Neurobiol Aging 96:12–21.

Weed MR, Taffe MA, Polis I, Roberts AC, Robbins TW, Koob GF, Bloom FE, Gold LH (1999) Performance norms for a rhesus monkey neuropsychological testing battery: Acquisition and long-term performance. Cogn Brain Res 8:185–201.

Yassa MA, Mattfeld AT, Stark SM, Stark CEL (2011) Age-related memory deficits linked to circuit-specific disruptions in the hippocampus. Proc Natl Acad Sci U S A 108:8873–8878.

